# Genome-wide analysis of the interplay between chromatin-associated RNA and 3D genome organization in human cells

**DOI:** 10.1101/2021.06.10.447969

**Authors:** Riccardo Calandrelli, Xingzhao Wen, John Lalith Charles Richard, Zhifei Luo, Tri C. Nguyen, Chien-Ju Chen, Zhijie Qi, Shuanghong Xue, Weizhong Chen, Zhangming Yan, Weixin Wu, Kathia Zaleta-Rivera, Rong Hu, Miao Yu, Yuchuan Wang, Wenbo Li, Jian Ma, Bing Ren, Sheng Zhong

**Author notes:** Equal contribution (John Lalith Charles Richard, Zhifei Luo). Correspondence: S.Z.

## Abstract

The interphase genome is dynamically organized in the nucleus and decorated with chromatin-associated RNA (caRNA). It remains unclear whether the genome architecture modulates the spatial distribution of caRNA and vice versa. Here, we generate a resource of genome-wide RNA-DNA and DNA-DNA contact maps in human cells. These maps reveal the chromosomal domains demarcated by locally transcribed RNA, hereafter termed RNA-defined chromosomal domains. Further, the spreading of caRNA is constrained by the boundaries of topologically associating domains (TADs), demonstrating the role of the 3D genome structure in modulating the spatial distribution of RNA. Conversely, stopping transcription or acute depletion of RNA induces thousands of chromatin loops genome-wide. Activation or suppression of the transcription of specific genes suppresses or creates chromatin loops straddling these genes. Deletion of a specific caRNA-producing genomic sequence promotes chromatin loops that straddle the interchromosomal target sequences of this caRNA. These data suggest a feedback loop where the 3D genome modulates the spatial distribution of RNA, which in turn affects the dynamic 3D genome organization.

## Introduction

The interphase genome is highly organized ^1^. The multiscale organizational features of the genome have been characterized, including A/B compartments^2^, topologically associating domains (TADs)^3,4^, and chromatin loops^5^. This multiscale organization begs the question of what the functions of such an intricate architecture are. Transcriptional regulation is one of the possible functions and the most extensively studied function. In this direction, the genome architecture is shown to regulate the transcription of specific genes ^1,6,7^, but it remains debatable whether the genome architecture has a widespread role in modulating the transcription of many genes ^8^. Moreover, it remains unclear if the 3D genome’s regulatory roles are limited to transcriptional regulation. Other possible functions have rarely been tested. Here, we test another possible function, namely regulating spatial localization of chromatin-associated RNA (caRNA) ^9^.

After initial debates ^9^, caRNA has been recognized as an integral component of interphase chromosomes rather than passive degradation products ^10–14^. Growing evidence confirms that caRNA regulates gene transcription and RNA splicing ^15–26^. These regulatory roles often depend on caRNA’s spatial localization within the nucleus ^20,27–30^. Depending on their spatial localizations, caRNA can orchestrate the organization of nuclear bodies and compartments ^29–32^ and foster the formation of transcriptionally silent or active chromosomal domains ^30,33,34^. However, it remains unclear how the caRNAs are spatially organized in the context of the multiscale genome architecture; whether there is any specificity in the spatial distribution of caRNAs; if there is, how is such specificity regulated; and in turn, whether the spatial localization of caRNA modulates the dynamic organization of the genome?

Guided by these questions, we generate high-resolution genome-wide RNA-DNA contact maps^15,35–38^ in human cells using *in situ* <u>M</u>apping of <u>R</u>NA-<u>G</u>enome Interaction (iMARGI)^35,36^. iMARGI captures RNA-genome associations by jointly sequencing caRNA and their associated genomic sequences with paired-end sequence reads^35^. iMARGI can differentiate the sequencing reads originating from RNA (iMARGI RNA-end reads) or genomic DNA (iMARGI DNA-end reads). We also use *in situ* Hi-C (Hi-C)^5,39^ to map genome-wide chromatin interactions. These maps reveal most caRNAs are associated with the genomic sequences within several megabases of their transcription sites.

To dissect any causal relationships between the 3D genome organization and caRNA, we generate RNA-DNA contact maps in the genetically engineered human cells where the TAD boundaries are deleted or inserted. Comparisons of these maps reveal the ability of TAD boundaries to constrain the spreading of caRNA on the chromosomes. These data demonstrate the 3D genome’s functions in regulating the spatial localization of the caRNA. Moreover, we generate RNA-DNA and DNA-DNA contact maps in human cells undergone either acute RNA depletion or deletion of a specific caRNA-producing sequence. These data reveal a suppressive role of between-anchor caRNA, i.e., the caRNA associated with the genomic region between the loop anchors, on chromatin looping. Thus, the spatial localization of the caRNA, in turn, modulates to the dynamic 3D genome organization.

## Results

### Localized RNA-genome association and RNA-defined chromatin domains

We generated iMARGI data from human embryonic stem (H1), foreskin fibroblast (HFF), and chronic myelogenous leukemia (K562) cells in duplicates (Table S1). These data revealed the relative level of any gene’s RNA attached to any genomic region (target region), hereafter called the RNA attachment level (RAL) of this gene and target region, defined as the number of iMARGI read pairs with the RNA ends mapped to this gene and the DNA ends mapped to this target region^35^. For example, in H1 ES cells the coding gene Jumonji and AT-Rich Interaction Domain Containing 2 (JARID2) exhibited large RAL in an approximately 5 Mb region containing the JARID2 gene (Figure 1a). Additionally, the non-coding gene Pvt1 Oncogene (PVT1) exhibited large RAL in an approximately 7 Mb region containing the PVT1 gene (Figure 1b). Overall, the average RAL of all the genes decreases as the genomic distance between the gene and the target region increases (Supplementary Figure S1a).

**Figure 1.**
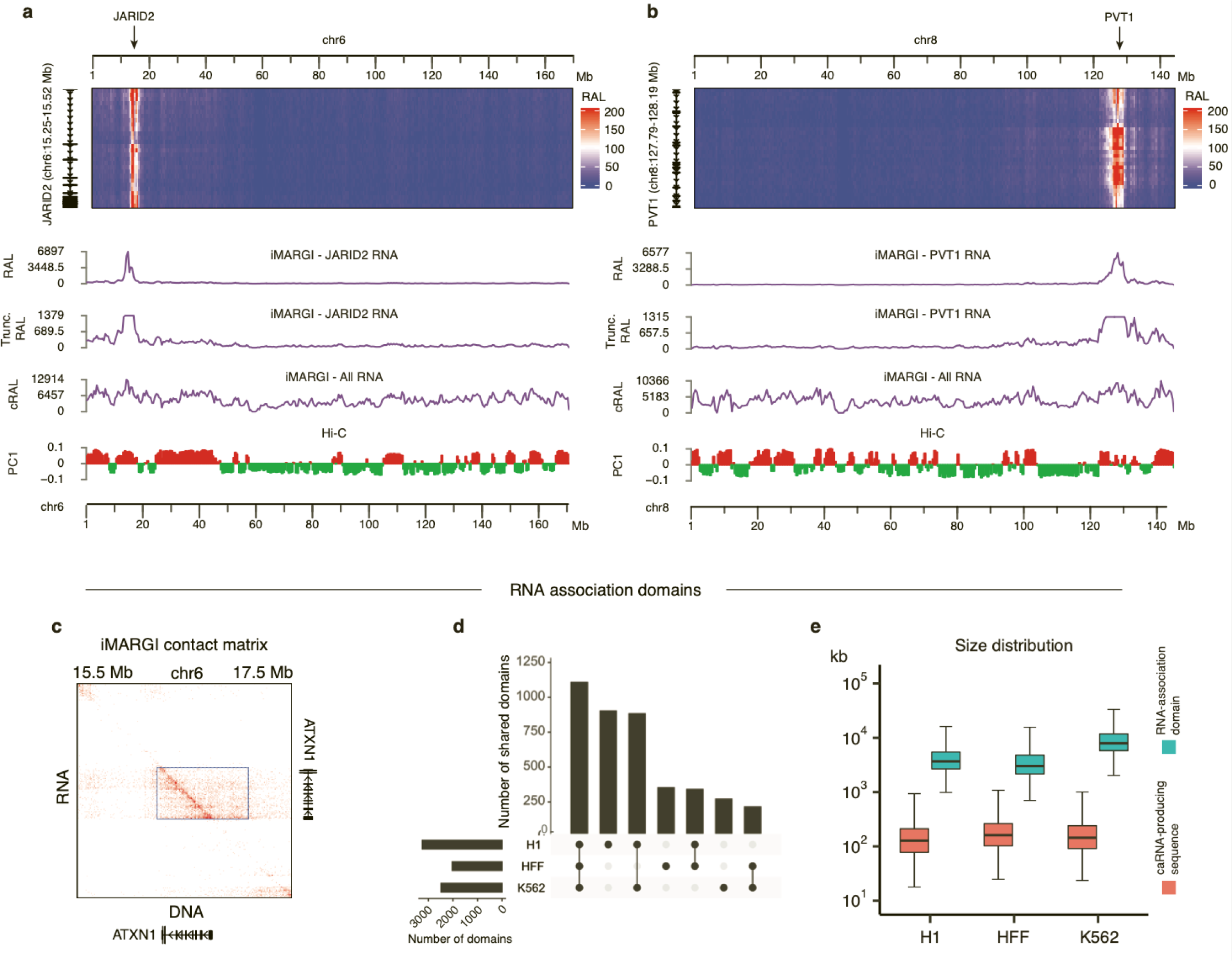
Localized RNA-genome association. (a) iMARGI’s contact matrix between the RNA of the JARID2 (rows, bin size = 10 kb) to the genomic sequence of chromosome 6 (columns, bin size = 500 kb). The RNA ciation level (RAL) of JARID2 RNA (RAL track), the truncated version of RAL showing the small values RAL track), and the cumulative RAL of all RNA (cRAL track) exhibit a correlation with the first principal onent of Hi-C’s contact matrix (PC1 track). (b) iMARGI’s contact matrix, RAL, and truncated RAL of the RNA on chromosome 6. (c) An RNA-DNA contact matrix in a 2M bp sequence on Chromosome 6. Each in this contact matrix represents the number of iMARGI read pairs with the RNA-end mapped to the sponding row and the DNA-end mapped to the corresponding column. The box marks an identified RNA-ciation domain, an approximately 1 Mb region containing the ATXN1 gene. (d) Upset plot of the numbers of etected RNA-association domains in H1, HFF, and K562. (e) Box plots of RNA-association domains’ sizes) corresponding to the widths of the detected rectangular blocks, and the lengths of caRNA-producing mic sequences (red) for each RNA-association domain corresponding to the heights of the detected ngular blocks in iMARGI’s contact matrix.

We represented iMARGI data as a contact matrix, where the rows represent the RNA ends of iMARGI read pairs, and the columns represent the corresponding DNA ends^36^ (Figure 1c). A notable difference to Hi-C’s symmetric contact matrix is that iMARGI’s contact matrix is asymmetric. This is because RNA-DNA contacts are not necessarily reciprocal. Rectangular blocks of high-value entries emerged as a recurring pattern from iMARGI’s contact matrix (Figure 1c). We identified the rectangular blocks using HOMER to call peaks on the rows of the contact matrix (row peaks), and in each row peak using HOMER to call one strongest peak in the columns (column peak). A pair of row peak and column peak defines a rectangular block. We identified 3,217, 2,019, and 2,468 rectangular blocks from H1, HFF, and K562 iMARGI data (Figure 1d). All the identified rectangular blocks overlap with the diagonal entries of iMARGI’s contact matrix, suggesting that they represent localized RNA-genome associations where a RNA’s target regions are near the transcription site of this caRNA. Each rectangular block corresponds to a unique chromatin domain, characterized by extensive genomic association of the RNA transcribed from within this domain. Hereafter we term such domains “RNA-association domains”. The size of an RNA-association domain, represented by the width of a rectangular block, can reach tens of megabases (Figure 1e). In summary, RNA-association domains emerged as a main feature of the genome-wide distributions of caRNA.

### Correlation between 3D genome compartmentalization and RNA-chromatin association

The 3D genome is organized on different scales, including compartments, TADs, and chromatin loops^1^. We asked whether the RNA association on any genomic region correlates with this genomic region’s 3D compartmentalization. To this end, we generated Hi-C data in H1, HFF, and K562 cells in duplicates and compared them with our iMARGI data (Table S1). We calculated the cumulative RAL (cRAL), the sum of the RAL of all the RNA, on every genomic region, defined as the number of iMARGI read pairs with the DNA ends mapped to this genomic region^35^. The A/B compartments as indicated by Hi-C contact matrix’s first eigenvector (PC1)^40^ exhibited a genome-wide correlation with cRAL (p-value < 2e-16, one way ANOVA), revealing a correlation between 3D genome compartmentalization and RNA-chromatin association.

We asked if the higher cRAL in the A compartment is completely attributable to a higher level of local transcription. To this end, we compared JARID2 and PVT1’s RALs with A/B compartments^40^ (PC1 track, Figure 1a, b). Both JARID2 and PVT1 exhibited small but non-zero RALs in several A compartment genomic regions that are tens of megabases away from the JARID2 and PVT1 genes (Figure 1a, b). However, the B compartment genomic regions that are closer to the JARID2 and PVT1 genes did not exhibit association of JARID2 or PVT1 RNA (Figure 1a, b), suggesting an enrichment of target regions of long-range RNA-chromatin contacts in the A compartment. Thus, the higher cRAL in the A compartment is not completely due to a higher level of local transcription.

### TAD boundaries insulate RNA-DNA contacts

TADs, where DNA sequences interact with each other more frequently than with the sequences outside, are important 3D genome features that are strongly correlated with transcriptional regulation^3,4^. We separately analyzed the RNA transcribed from within a TAD or the other regions of the same chromosome outside of this TAD. The chromatin attachment level of any RNA transcribed from within a TAD sharply decreases at the two boundaries of this TAD (p-value = 6.5e-16, Wilcoxon rank-sum test) (Figure 2a). Conversely, the attachment level of any RNA transcribed from outside of a TAD exhibits drastic changes at the TAD boundaries in the opposite direction (p-value = 2.6e-12, Wilcoxon rank-sum test) (Supplementary Figure S1b). These changes at TAD boundaries cannot be completely explained by the 1-dimensional genomic distance to the caRNA’s transcription site. They suggest the possibility that a TAD boundary can insulate RNA-DNA contacts from the two sides of this boundary (cross-over RNA-DNA contacts).

**Figure 2.**
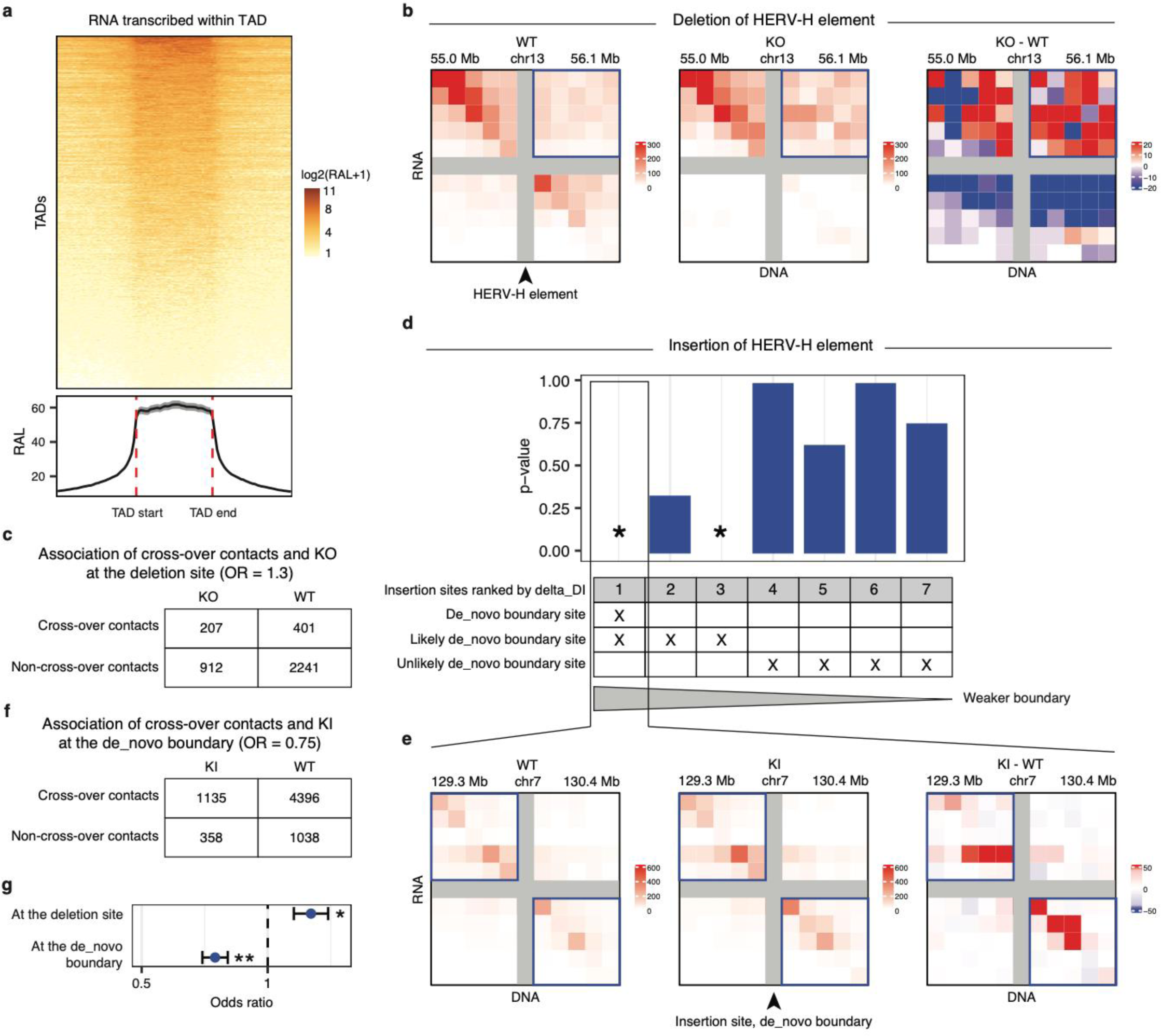
TAD boundaries suppress cross-over RNA-DNA contacts. (a) The RNA association level (RAL, color-d) of the RNA transcribed within each TAD (row) on this TAD (center block) and its equal-length flanking ns (x axis). Curve at the bottom: the average RAL of all TADs (rows). (b) Comparison of normalized RNA-contact matrices in WT and KO cell lines. The arrowhead points to the HERV-H element in WT that is ed in KO. KO-WT: The contrast of the KO and WT contact matrices where red indicates an increase of DNA contacts in KO. The increased RNA-DNA contacts in KO are enriched with cross-over contacts (in ox at the upper right corner). (c) The 2×2 contingency table for an association test based on the data in l b. (d) The seven previously identified insertion sites (columns) are ranked by delta_DI, where a smaller _DI (on the right) indicates a smaller increase in the ability to insulate cross-over DNA-DNA contacts (a er putative boundary). The rows mark whether each insertion site is a “de_novo boundary site” (Column 1), ely-de_novo-boundary site” (Columns 1-3), or an “unlikely-de_novo-boundary site” (Columns 4-7), based e comparison of Hi-C data in KI and WT. A Chi-square test is performed on each insertion site (column) d on the iMARGI data in KI and WT. A smaller p-value (y axis) represents stronger evidence against the ypothesis that there is no association between the RNA-DNA cross-over contacts and KI. *: p-value < 1.0e-(e) Comparison of normalized RNA-DNA contact matrices in WT and KI cell lines at the de_novo boundary arrowhead). KI-WT: The contrast of the KI and WT contact matrices where red indicates an increase of DNA contacts in KI. The increased RNA-DNA contacts in KI are enriched in the non-crossover contacts (in oxes at the upper left and lower right corners). (f) The 2×2 contingency table for an association test based e data in panel e. (g) Significance levels for the deletion site (panel b, c) and the de_novo-boundary insertion panel e, f). *: p-value < 0.05, **: p-value < 1.0e-4.

We asked if altering the genomic sequence within a TAD boundary can affect the cross-over RNA-DNA contacts. First, we leveraged our previous finding that a CRISPR-mediated deletion of a HERV-H element (Chr13:55,578,227-55,584,087) (KO) within a TAD boundary from H9 human ES cells (WT) abolishes this TAD boundary^41^. We carried out iMARGI experiments on the KO and WT cells. We counted the numbers of cross-over and non-cross-over iMARGI read pairs in WT and KO. Compared to WT, KO exhibited an increased proportion in the cross-over read pairs (OR = 1.3, p-value = 0.013, Chi-square test) (Figure 2b, c, g). Thus, deleting a fraction of a TAD boundary reduced its insulation to cross-over RNA-DNA contacts.

Second, we previously created an insertion cell line (KI) using piggyBac transposon-mediated genomic insertion of this HERV-H sequence and identified seven insertion sites in KI^41^ (Columns, Figure 2d). Four of the seven insertion sites exhibited small increases in insulation (unlikely-de_novo-boundary sites), as measured by the difference in directionality index (delta_DI < 20) (Columns 4-7, Figure 2d), whereas the other three insertion site exhibited large increases in insulation (delta_DI > 20, likely-de_novo-boundary sites)^41^ (Columns 1-3, Figure 2d). Only one insertion site, that has the largest increase in insulation (delta_DI = 66.3), reached the significance level to be detected as a *de novo* TAD boundary, i.e., a boundary called in the KI Hi-C but not called in the WT Hi-C (de_novo-boundary site) (Column 1, Figure 2d). We note the de_novo-boundary site is one of the three likely-de_novo-boundary sites.

To test any impact of any insertion site on RNA-DNA contacts, we carried out iMARGI in KI and WT cells. For every insertion site, our null hypothesis is that whether any RNA-DNA contact is a cross-over or a non-cross-over contact is independent of whether this RNA-DNA contact is detected in KI or WT. The three likely-de_novo-boundary sites (delta_DI > 20) all led to some degrees of decrease in the odds ratio (OR) of the cross-over RNA-DNA contacts in KI (OR < 0.90), in which the decreases on two of the three likely-de_novo-boundary sites were significant (p-value < 1.0e-4, Chi-square test, stars in Figure 2d). In particular, the de_novo-boundary site exhibited a significant decrease in the OR of the cross-over RNA-DNA contacts in KI (OR = 0.75, p-value = 3.7e-5, Chi-square test, Figure 2e-g). Thus, the *de novo* creation of a TAD boundary suppressed cross-over RNA-DNA contacts.

In contrast, none of the four unlikely-de_novo-boundary sites (delta_DI < 20) led to a detectable decrease in the OR of the cross-over RNA-DNA contacts in KI (OR > 0.99, p-value > 0.63, Chi-square test, Figure 2d). Thus, inserting the same DNA sequence without sufficient subsequent changes in TAD structure did not suppress the cross-over RNA-DNA contacts. Taken together, these data show an impact of the 3D genome structure to the distribution of caRNA.

### Induction of transcription locally suppresses chromatin looping

Our next question is whether RNA has an impact on the 3D structure of the genome. We approached this question in three steps. First, we depleted RPB1, the largest subunit of RNA Polymerase II (RNAP II), in HCT116 cells. RPB1 depletion resulted in increases in loop number and strengths as measured by Hi-C (Extended Text: RPB1 depletion). This result is consistent with the recent report in another cell line (DLD-1), where depletion of RPB1 led to the emergence of chromatin loops^42^. These data implicate the transcriptional machinery, especially the presence of RNAPII on chromatin in suppressing chromatin looping.

Second, we tested if the induction of the transcription of a specific gene suppresses chromatin looping. To this end, we leveraged that there is a ∼55 kb loop straddling the AAVS1 locus (AAVS1 loop), and nearby, there is a non-overlapping loop with a similar size to the AAVS1 loop^43^ (Nearby Ctrl loop, Figure 3a). We applied doxycycline to engineered H1 ES cells with a dCas9-KRAB knock-in transgene at the AAVS1 locus^44^ to induce transcription of the transgene at the AAVS1 locus (Dox+), and subsequently tested the loops with chromosome conformation capture (3C)^45^. Compared to without doxycycline (Dox-), Dox+ weakened the AAVS1 loop (star, Figure 3b) but had little impact on the Nearby Ctrl loop (Figure 3b). Thus, inducing the expression of a gene can suppress a chromatin loop that straddle across this gene. Taken together, the transcription of a gene can locally suppress chromatin looping. It remains unclear whether it is the transcriptional machinery, the process of transcription, or the product of transcription, i.e., RNA, that affects chromatin looping.

**Figure 3.**
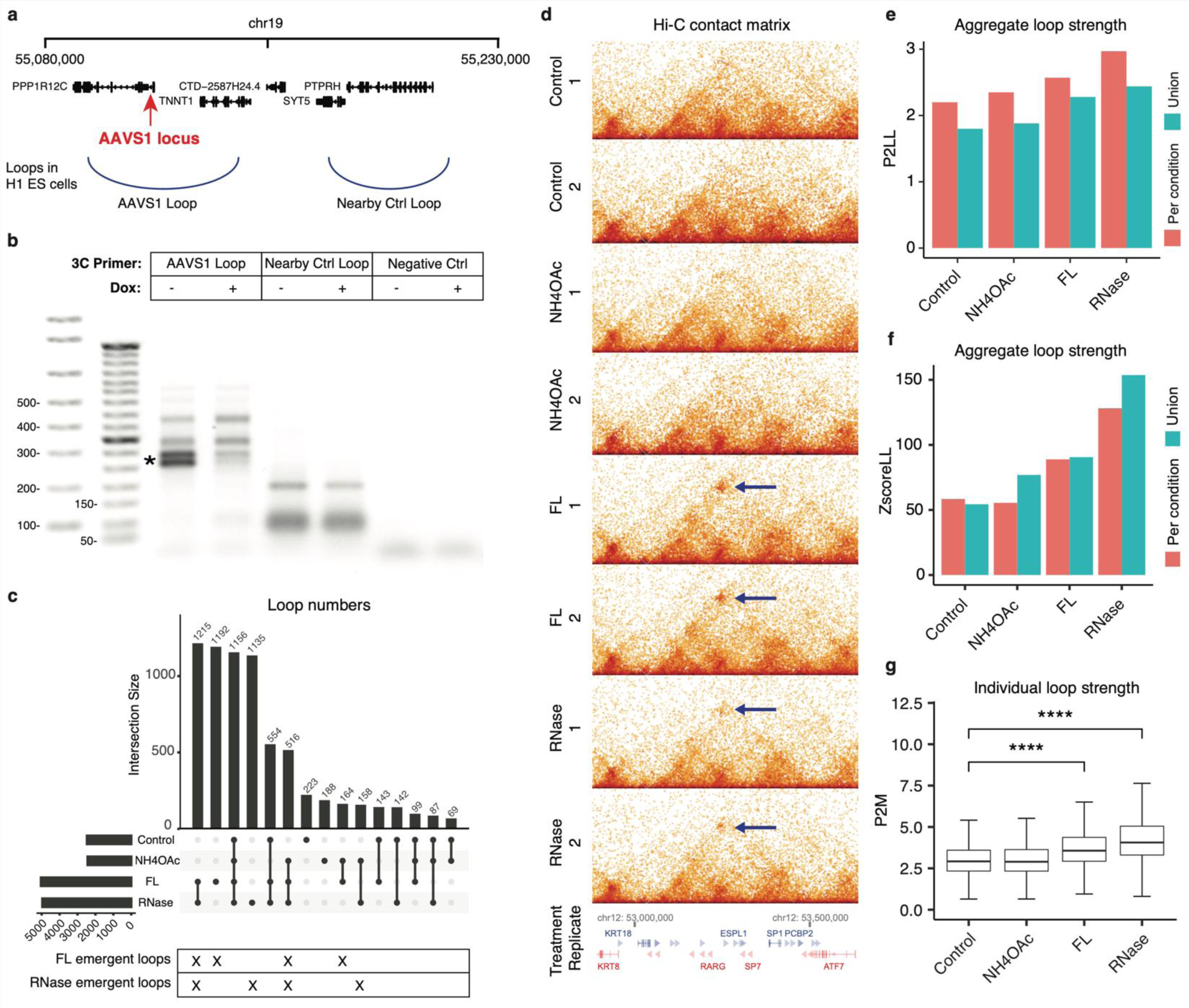
RNA-related loop changes. (a) Transcription induction of a gene suppresses a loop straddling this gene. mic coordinates of the AAVS1 locus, the loop straddling the AAVS1 locus (AAVS1 loop), and a nearby loop similar size (Nearby Ctrl loop). (b) 3C products without doxycycline (Dox: -) and with transcription induction xycycline (Dox: +), based on primers against the AAVS1 loop (Lanes 3, 4), the Nearby Ctrl loop (Lanes 5, nd a size-matched control region without any Hi-C detected loop (Negative Ctrl, Lanes 7, 8). *: Difference products between Dox- and Dox+. Lane 1: E-Gel™ 1 kb DNA Ladder. Lanes 2 and 9: E-Gel™ 50 bp DNA er. (c) Upset plot of the loop numbers in the four conditions, control, NH4OAc, FL, and RNase (rows). (d) ample of loop changes. Hi-C contact matrix of every replicate (row). Arrows: a shared loop in FL and RNase s absent in control and NH4OAc. (e-g) FL and RNase increase loop strengths. (e, f) Aggregate loop strength sented by P2LL (e) or ZscoreLL (f) (y axis) in each condition (column). Color bars: the loops detected in condition (red) or their union (blue). (g) Box plots of the strengths of individual loops (P2M) in every tion (column). ****: p-value < 2.2e-16.

### RNA has a genome-wide impact on chromatin looping

We asked whether it is the transcription (including the association of the transcriptional machinery on chromatin and the process of transcription) or the RNA that impacts chromatin looping. We recognized that we could not answer this question by only testing with a specific genomic locus. This is because the answer at one genomic locus cannot necessarily rule out the alternative answer in other genomic regions. Thus, we recognized that the perhaps more important question is whether transcription or RNA has a genome-wide impact on chromatin looping. Furthermore, we recognized that if RNA impacts chromatin looping, then the transcriptional process must be implicated. However, if the transcriptional process is the cause, the causal chain does not necessarily involve RNA. With these considerations, we revised our question to whether RNA can impact chromatin looping genome-wide? To answer this question, we compared chromatin looping in control, transcription-inhibited ^46–48^, and RNase-treated cells ^49^. If the primary cause is the transcription process, we expect to see a widespread impact in transcription inhibition but not in acute RNase treatment. However, if transcription inhibition and acute RNase treatment lead to overlapping changes in chromatin loops genome-wide, the data would suggest RNA is involved in modulating chromatin looping. Additionally, we included another experimental condition where electrostatic molecular interactions are inhibited to test if any observed impacts are attributable to charge-driven condensates or phase separation^50,51^.

In our third step, we subjected H1 cells with ammonium acetate (NH4OAc) to disrupt electrostatic molecular interactions (the interactions due to electric charges) ^50–52^, flavopiridol (FL) to suppress transcription elongation without displacing RNAPII from chromatin ^46–48 53^, and acute RNase treatment to reduce RNA in the nuclei (10-minute RNase treatment before fixing the cells) ^49^ based on established protocols (NH4Oac ^50^, FL ^46^, RNase A ^49^). NH4OAc disrupts molecular electrostatic interactions in living cells by providing monovalent cations without perturbing intracellular pH^51^. To check the expected effects of the three treatments, we immunostained nuclear speckle-associated proteins SON^54^ and SC35^55^ in control and each treatment. NH4OAc reduced the numbers of SON and SC35’s foci (p-value = 0.001 for SON, 0.009 for SC35, Wilcoxon test) (Supplementary Figure S2a, b, e, f, i, l), consistent with the role of RNA’s electrostatic interactions in maintaining nuclear speckles^56^. Conversely, FL made SON and SC35 foci larger and more distinct^57^ (Supplementary Figure S2c, g, j, m). RNase A increased the numbers of SON and SC35’s foci (p-value = 0.034 for SON, 0.010 for SC35, Wilcoxon test) (Supplementary Figure S2d, h, k, n), consistent with the observations that “low RNA/protein ratios promote phase separation into liquid droplets”^58^ and condensate formation^59^.

We generated Hi-C after each treatment in duplicates (Table S1) and analyzed these data together with those of the unperturbed H1 cells (control). We called chromatin loops from our Hi-C data in each of the four conditions that have comparable sequencing depths (Table S1) using HiCCUPS^60^. The loop numbers were similar in control (2,473 loops) and NH4OAc (2,437 loops) (p-value = 0.55, paired t test) and were increased in FL (5,039 loops) (p-value < 1.1e-8, paired t test) and RNase (4,963 loops) (p-value < 2.3e-9, paired t test) (Figure 3c). These loop number differences cannot be attributed to different sequencing depths or batch effects because the samples were prepared in the same batch and sequenced to comparable depths (600 – 650 million read pairs per condition, Table S1b). Most of the emerged loops in FL colocalized with the emerged loops in RNase (first column, Figure 3c). For example, a loop linking ATF7 and KRT18 genes that was absent in control and NH4OAc emerged in both FL and RNase (arrows, Figure 3d, Supplementary Figure S3).

The overall loop strength was similar in control and NH4OAc, but stronger in FL and RNase, as reflected by both Peak to Lower Left (P2LL) (Figure 3e) and Z-score Lower Left (ZscoreLL) scores^5^ (Figure 3f). We repeated these analyses based on the union of the loops in the four conditions and quantified every loop’s strength by Peak to Mean (P2M) in each condition. P2Ms were greater in FL and RNase than in control (p-value < 2.2e-16, Wilcoxon test), whereas NH4OAc’s P2Ms were not different from the control’s (p-value = 0.41, Wilcoxon test) (Figure 3g). The consistent increases of loop strengths in FL and RNase as compared to control support our detected increases of loop numbers in FL and RNase and suggest that our conclusion of loop number increases does not depend on the threshold of loop calls. Taken together, FL and RNase both resulted in an increase of chromatin loops and these emerged loops often co-localize. As opposed to the null hypothesis that RNA does not have a genome-wide impact on chromatin looping, these data are in favor of a suppressive effect of RNA to chromatin looping genome-wide.

### RNA’s genomic target regions correlate with the suppressed chromatin loops

Our next question is what RNA has an impact on which chromatin loops. Although we cannot analyze every aspect of a select RNA, we can analyze the chromatin-associated fraction of this RNA, in terms of this RNA’s genomic target regions (target region) and the RNA attachment level (RAL) of this RNA on any target region. We can compare the target region with the genomic location of any chromatin loop. Thus, we asked whether the change of any caRNA, in terms of changes of target regions or RAL, correlates with the change of any chromatin loop. Answering this question can inform us which RNA could impact which chromatin loop, although we will miss those impacts that are independent of RNA-chromatin association.

We generated iMARGI in each treatment condition (NH4OAc, FL, RNase) in duplicates (Table S1) and analyzed these data together with those of the unperturbed H1 cells (control). As expected, FL exhibited the largest reduction of the heights of the rectangular blocks in iMARGI’s contact matrix (p-value < 3e-104, Wilcoxon rank-sum test) (Supplementary Figure S4e), consistent with FL’s inhibitory effect on transcription elongation^47^. RNase exhibited the largest reduction of caRNA domains’ number (3,217 in control and 357 in RNase, p-value < 3e-9, paired t test) (Supplementary Figure S4d) and sizes (widths of the rectangular blocks) (p-value < 5e-210, Wilcoxon rank-sum test) (Supplementary Figure S4f).

We analyzed two groups of caRNA, namely those associated with loop anchors (anchor caRNA), and those between loop anchors (between-anchor caRNA). We asked if the changes in the level of between-anchor caRNA correlates with the changes of chromatin loops across our treatment conditions. To answer this question, we analyzed the union of the loops (Union loops) detected in every condition (Control, NH4OAc, FL, RNase). These Union loops represent all possible loop locations, including those detected as loops in the Control (control loop) or in an RNA perturbation experiment (emergent loop). We used the ratio of between-anchor caRNA and anchor caRNA levels (Inside-loop To Anchor ratio (ITA ratio)) to represent the relative level of between-anchor caRNA for any Union loop.

First, we tested whether the detected loops in control (control loops) tend to locate at the genomic locations with a low level of between-anchor caRNA in the control. We carried out this test using Gene Set Enrichment Analysis (GSEA)^61^. According to GSEA’s procedure, we sorted the Union Loops by increasing levels of between-anchor caRNA, i.e., increasing ITA ratios, creating a ranked list (Figure 4a). We then plotted the corresponding GSEA score at every rank (Figure 4b), where a positive/negative GSEA score indicates an enrichment/depletion of the control loops in the subset of top-ranked Union Loops. Here, top-ranked means from rank #1 to the current rank on which GSEA score is reported. The GSEA scores stayed positive in the top portion (∼30%) of this rank list (Figure 4b), suggesting that the control loops are enriched in those Union loops that exhibit lower levels of between-anchor caRNA in the control condition than in the other conditions. In other words, among all the locations where loops have been detected, the control loops tend to appear at those locations where the relative level of between-anchor caRNA is low.

**Figure 4.**
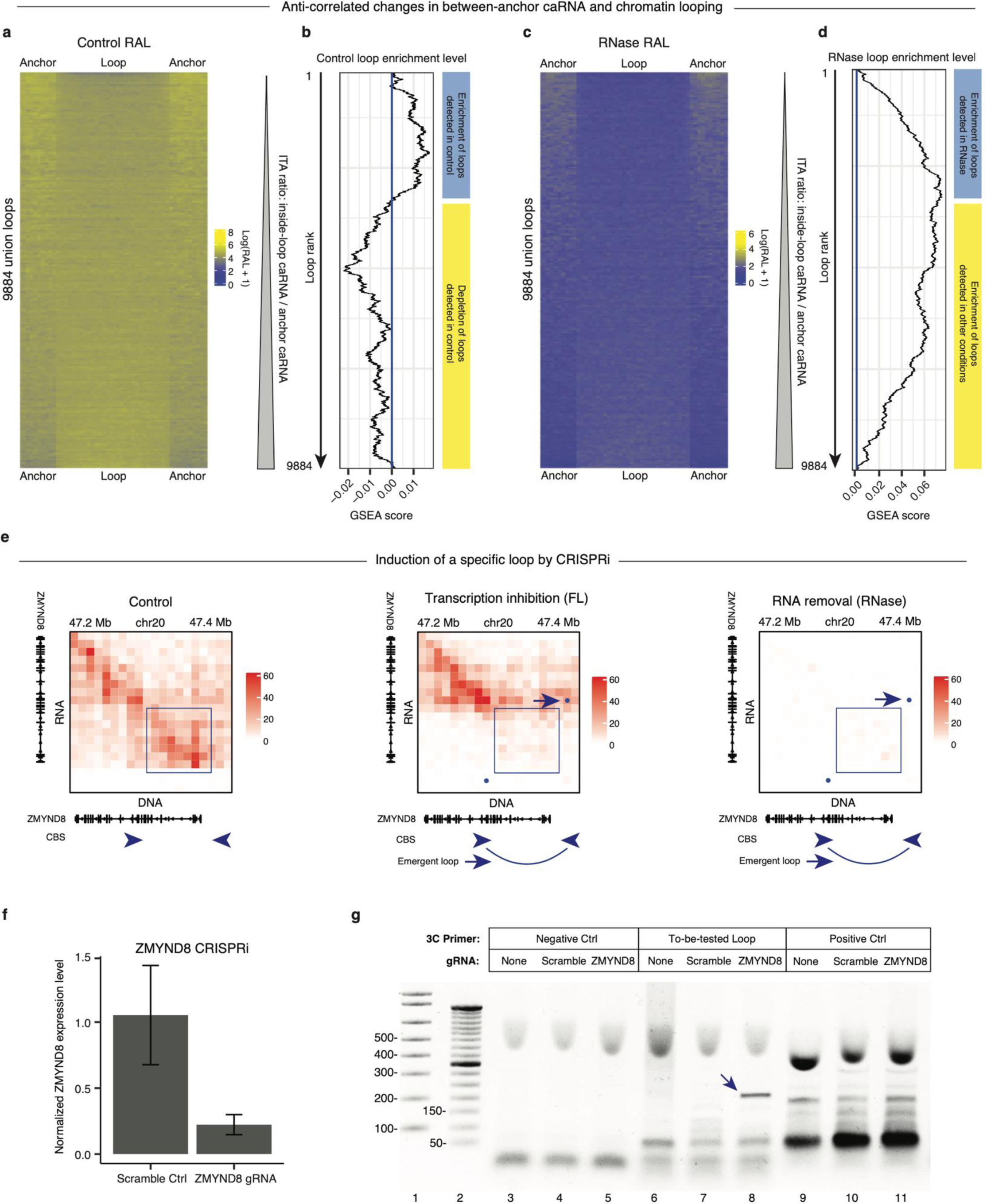
Between-anchor caRNA anticorrelates with chromatin looping. (a-b) The loops in the Control (control) are depleted with between-anchor caRNA. (a) The caRNA levels in the control (Control RAL) on loop ors (two sides) and between the anchors (middle) is color-coded (blue: low, yellow: high) for every loop ted in any condition (Union loops, rows). Loops are ranked by the relative level of their between-anchor A (Inside-loop To Anchor (ITA) ratio) from low (top) to high (bottom). (b) The enrichment/depletion level A score, x axis) of the control loops in the subset of loops from the top-ranked loop (first row) and the ntly ranked loop (current row, y axis). A positive/negative GSEA score indicates an enrichment/depletion of ontrol loops in this subset of loops. The control loops are enriched in the top-ranked loops, i.e., those with vels of between-anchor caRNA (blue bar on the right), and are depleted in the bottom-ranked loops, i.e., with high levels of between-anchor caRNA (yellow bar). (c-d) RNase emergent loops are those with low of between-anchor caRNA. (c) The caRNA levels in the RNase (RNase RAL) on loop anchors (two sides) etween the anchors (middle) is color-coded (blue: low, yellow: high) for every loop detected in any condition n loops, rows). The union loops (rows) are ordered by the relative level of their between-anchor caRNA TA ratio calculated in RNase) from low (top) to high (bottom). (d) The enrichment level (GSEA score, x axis) RNase loops (x axis) in the subset of loops from the top-ranked loop (first row) and the currently ranked (current row, y axis). The RNase-specific loops are enriched in the top-ranked loops, i.e., the loops with low of between-anchor caRNA in RNase, as indicated by the increasing GSEA scores (blue bar on the right). ntrast, the loops detected in other conditions are enriched in the bottom-ranked loops, i.e., the loops with evels of between-anchor caRNA in RNase, as indicated by the decreasing GSEA scores (yellow bar). (e-anscriptional suppression of the ZMYND8 induces a specific chromatin loop. (e) Changes in iMARGI RNA-contact maps in Control (left panel), FL (central panel), and RNase (right panel). FL reduced the caRNA in pstream region of the ZMYND8 gene (blue dox) and induced a chromatin loop near the caRNA-depleted n (curve at the bottom). RNase reduced the caRNA from a wider genomic region and induced the same matin loop as that in FL. CBS: CTCF binding site. Arrowheads point to CBSs’ directions. Blue dots: Hi-C ed loops that are superimposed on this iMARGI contact map. Arrow: the emergent loop in FL. (f) parison of normalized ZMYND8’s expression levels (y axis) in CRISPRi experiments with the scrambled A (Scramble Ctrl) and ZMYND8-targeting gRNA (ZMYND8 gRNA). (g) 3C products from the Negative Ctrl rs (the first 3 lanes), the primers for the To-be-tested loop (3 middle 3 lanes), and the Positive Ctrl primers ast 3 lanes) in CRISPRi experiments without a gRNA (gRNA: None), with a scrambled gRNA control (gRNA: mble), or with the ZMYND8-targeting gRNA (gRNA: ZMYND8). The Negative Ctrl primers did not yield any ct in any experiment (Lanes 3-5). The Positive Ctrl primers yielded products of the same sizes in all three riments (Lanes 9-11). The primers for the to-be-tested loop yielded a product with ZMYND8-targeting gRNA w) but not with a scrambled gRNA or without gRNA (Lanes 6-8), confirming that a loop is created by ND8 CRISPRi. Lane 1: E-Gel™ 1 kb DNA Ladder. Lanes 2: E-Gel™ 50 bp DNA Ladder.

Second, RNase reduced caRNA levels in all Union loops (Figure 4c) and nearly doubled the number of detected loops as compared to Control (Figure 3c). We tested whether the loops in RNase (RNase loops) appeared at the locations where the between-anchor caRNA is most rigorously depleted in RNase. To this end, we re-ordered the Union loops by increasing levels of between-anchor caRNA, i.e., the ITA ratio in RNase (Figure 4c). As expected, all the GSEA scores are positive in this analysis (Figure 4d), which is because the RNase loops comprise the majority (∼75%) of the Union loops, and therefore a majority in any top ranked subset. The GSEA scores in this rank list of Union loops first increased and then decreased, which means that the RNase loops are enriched in the higher ranked subset, which are the Union loops with low levels of between-anchor caRNA in RNase. This enrichment means that the emerged loops in RNase often appeared at the genomic locations where the between-anchor caRNA is most rigorously removed by RNase. Taken together, we observed a genome-wide negative correlation between between-anchor caRNA and the chromatin loops that stride across the target region of these caRNA.

### Reducing select RNA creates specific chromatin loops

We wondered if we could apply the aforementioned correlation to identify which RNA has an impact on what chromatin loops. To this end, we tested whether we could create a particular chromatin loop by reducing a specific RNA, which is the RNA that exhibits a strong level of chromatin attachment to the genomic region between the anchors of this chromatin loop. We chose the ZMYND8 RNA for this test. We chose the ZMYND8 RNA because (1) FL reduced the RAL of the ZMYND8 RNA in an approximately 90 kb genomic region (Figure 4e); (2) RNase also removed the ZMYND8 caRNA in this (and a larger) genomic region; (3) a chromatin loop straddling across this 90 kb region was detected by Hi-C in both FL and RNase (arrows, Figure 4e), hereafter called the “straddling loop”. We note that FL and RNase reduce the RALs of many RNAs, and thus from these data we cannot conclude that the emergence of this straddling loop in FL and RNase is due to the reduction of any specific RNA.

RNA knockdown without affecting transcription can reduce nucleoplasmic RNA and suppress long-range RNA-chromatin interactions, however, it cannot effectively remove nascent RNA that are associated with the chromatin near the transcription locus ^21^. Therefore, we do not have a method to effectively remove ZMYND8 caRNA near the ZMYND8 gene without affecting the transcription of the ZMYND8 gene. We employed two approaches to address this issue. First, we asked whether suppression of ZMYND8 transcription has the same effect as RNase in creating the “straddling loop”. Second, we will describe in subsequent sections the analysis of inter-chromosomal RNA-chromatin interactions, where we can better distinguish between impacts of the RNA from the transcriptional process.

We suppressed ZMYND8 by CRISPR interference (CRISPRi) in an H1 ES cell line with doxycycline-inducible dCas9-KRAB^44,62^. Compared to scrambled gRNA control, our gRNA targeting ZMYND8’s promoter reduced ZMYND8’s transcription level to approximately 25% (Figure 4f). We designed chromosome conformation capture (3C) primers^45^ for (1) a negative control “loop” (Negative Ctrl) that is located 200 kb upstream of the emerged loop and has approximately the same size as the emerged loop, which is not detected as a loop in any Hi-C experiment, (2) a positive control loop (Positive Ctrl) detected by Hi-C in both control and FL, which is not on the same chromosome as ZMYND8, and (3) the straddling loop (also termed the “to-be-tested loop”). We carried out 3C after treating the cells with doxycycline without supplying gRNA (gRNA:None Ctrl), supplying with a scrambled gRNA (gRNA:Scramble Ctrl), and with gRNA targeting ZMYND8’s promoter (gRNA:ZMYND8). The Negative Ctrl primers did not yield any product in any experiment (the first 3 lanes), and the Positive Ctrl primers yielded products at the expected sizes in all three experiments (the last 3 lanes, Figure 4g). In contrast, the primers for the to-be-tested loop yielded a unique product with ZMYND8 gRNA (arrow, Figure 4g), which is absent from the gRNA:None and gRNA:Scramble controls. In summary, acute reduction of RNA induced many chromatin loops including the straddling loop, and suppression of the ZMYND8 expression can re-create the emergence of the straddling loop. Thus, the negative correlation of between-anchor caRNA and chromatin loops can help to identify which RNA has an impact on which chromatin loop. We note that the CRISPRi experiment by itself cannot distinguish whether the loop was created by suppression of transcription or reduction of ZMYND8 RNA. This CRISPRi experiment demonstrates that a loop created by acute depletion of RNA (RNase) can be re-created by suppression of the expression of a specific gene.

### Acute RNA reduction increases the average strength of the loops with convergent CTCF binding sites in their loop anchors

Our next question is whether an RNA can impact the chromatin loops located far from the transcription locus of this RNA (distal loops). We recognize that our previously mentioned correlation is not sufficient to connect a specific RNA to specific distal loops. This is because an RNA can associate with many distal genomic regions, often at low levels. Thus, we proceeded to identify additional correlational rule(s) to between RNA and chromatin loops.

Convergent CTCF binding sites (CBS) in the loop anchors is a characteristic of the loops created by loop extrusion^59^. We tested whether the convergent CBS are enriched in the anchors of the loops with increased loop strengths in RNase. To this end, we categorized the Union Loops (the union of the loops detected in any treatment condition) into three groups based on the orientations of the CTCF binding sites at their anchors, namely the loops with convergent CBS, non-convergent CBS, or no CBS. We used Peak to Lower Left (P2LL) to quantify the strength of each loop^5^. Compared to control, RNase treatment increased P2LL in the Union Loops with convergent CBS (p-value < 1.6E-9, Wilcoxon test, Figure 5a). In comparison, RNase did not increase P2LL in the Union Loops with non-convergent CBS (p-value = 0.4663, Wilcoxon test, Figure 5a) or in the loops without CBS (p-value = 0.6277, Wilcoxon test, Figure 5a). Thus, acute RNA reduction increased the average strength of those loops with convergent CBS in their loop anchors, suggesting an enrichment of convergent CBS in anchors of RNA suppressed loops. In summary, we have observed two genome-wide correlations, which are (1) a negative correlation of between-anchor caRNA and chromatin loops and (2) an enrichment of convergent CBS in RNA suppressed loops. Hereafter we call these correlations the “correlational rules”.

**Figure 5.**
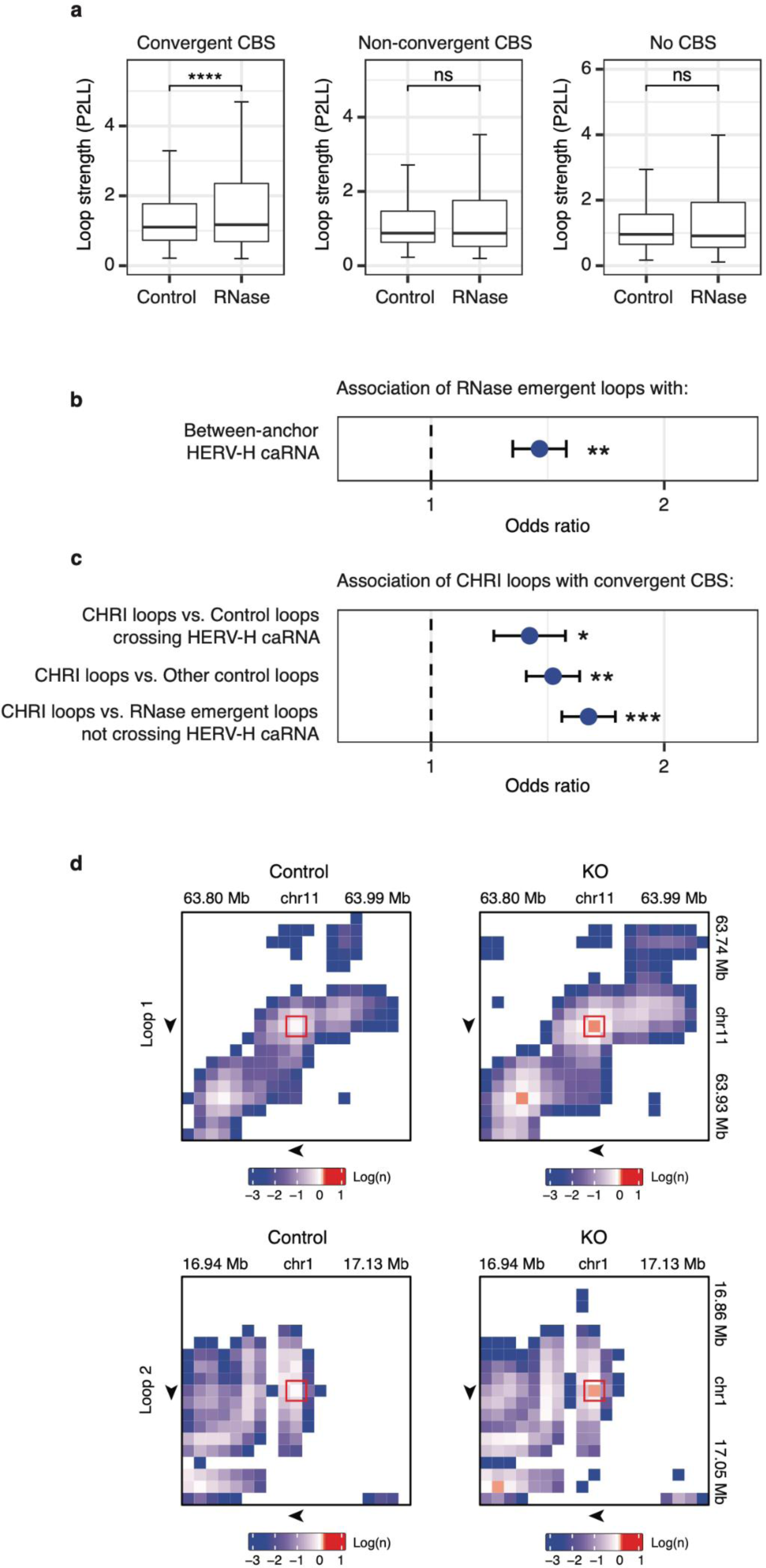
Enrichment of convergent CTCF binding sites (CBS) in the anchors of RNA-affected loops. (a) parison of loop strengths (P2LL, y axis) in the loops with convergent CBS, non-convergent CBS, and without in the control and RNase (columns). ****: p-value < 1.6e-9. ns: not significant. (b) Enrichment of RNase gent loops with between-anchor HERV-H caRNA in control (odds ratio, x axis). **: p-value < 5.6e-5, Chi-re test. Odds ratio > 1 means enrichment. (c) Enrichment of “candidate HERV-H caRNA insulated loops” I-loops) with convergent CBS in their loop anchors as compared to control loops striding across HERV-H A-attached genomic sequences (first row), the other control loops not striding across HERV-H caRNA-hed genomic sequences (second row), and the RNase emergent loops not striding across any HERV-H A-attached genomic sequences (third row). *: p < 6.8e-3, **: p < 5.8e-6, ***: p < 4.1e-9. (d) The two target-ing loops (rows) with increased Hi-C contacts in HERV-H KO (KO column) as compared to control (Control n). Hi-C data were denoised using the DeepLoop software. The denoised Hi-C contact maps were shown log scale. Arrow: direction of CTCF binding site in the loop anchor.

### Removal of select RNA increases the strengths of a subset of distal chromatin loops

We wondered if we could apply the correlational rules to identify which RNA may have an impact on what distal chromatin loops. To this end, we tested whether removing specific RNA can increase the strengths of certain distal loops. We chose the HERV-H RNA for this test for the following reasons. First, we identified the HERV-H caRNA-associated genomic sequences (HERV-RNA target regions) in Control and compared them with the locations of the loops emerged in RNase (RNase emergent loops). The RNase emergent loops are enriched at the locations that exhibit between-anchor HERV-H caRNA in Control (odds ratio = 1.38, p-value = 5.621e-5, Chi-square test, Figure 5b), suggesting that those loops that stride across between-anchor HERV-H caRNA are suppressed in Control. Second, we analyzed the subset of RNase emergent loops that stride across HERV-H caRNA-attached genomic sequences in Control. Hereafter, we call this subset of RNase emergent loops as “<u>c</u>andidate <u>H</u>ERV-H ca<u>R</u>NA insulated loops” (CHRI-loops). CHRI-loops are enriched with convergent CBS in their loop anchors as compared to (1) control loops striding across HERV-H caRNA-attached genomic sequences (OR = 1.34, p = 0.0068, Chi-square test), and to (2) the other control loops not striding across HERV-H caRNA-attached genomic sequences (OR = 1.44, p = 5.8e-6, Chi-square test), and to (3) the RNase emergent loops not striding across any HERV-H caRNA-attached genomic sequences in Control (OR = 1.60, p = 4.1e-9, Chi-square test, Figure 5c). Thus, convergent CBS are enriched in the loop anchors of CHRI-loops.

We tested whether deleting an HERV-H element from the human genome can lead to increase the loop strength of any distal CHRI-loop. To this end, we re-used our Chr13:55.5MB_HERV KO (CRISPR-mediated deletion of a HERV-H element at Chr13:55,578,227-55,584,087) human ES cells ^41^. We identified the caRNA transcribed from this Chr13:55.5MB_HERV element and its target genomic sequences (Chr13:55.5MB_HERV targets) in the WT. We call the loops that stride across any Chr13:55.5MB_HERV targets as “target-crossing loops”. We compared the loop strength changes of all target-crossing loops between Chr13:55.5MB_HERV KO and WT based on Hi-C data. No target-crossing loop exhibited detectable decrease in loop strength in Chr13:55.5MB_HERV KO, whereas two target-crossing loops exhibited increased loop strengths in the Chr13:55.5MB_HERV KO (Figure 5d). Both Chr13:55.5MB_HERV KO-induced target-crossing loops contain convergent CBS in their loop anchors (Figure 5d). Neither Chr13:55.5MB_HERV KO-induced target-crossing loop locates on Chromosome 13, where the HERV-H element is deleted (Figure 5d). Thus, removal of specific RNA increased the loop strengths of a subset of chromatin loops that stride across this RNA’s interchromosomal target regions and contain convergent CBS in their loop anchors. These data suggest specific RNA can modulate a subset of chromatin loops. Furthermore, the correlational rules help to identify which RNA modulates what chromatin loops.

## Discussion

We presented a resource composed of genome-wide RNA-DNA and DNA-DNA contact maps in three human cell lines. The iMARGI and Hi-C experimental protocols and data processing pipelines used for generating this resource were proven by the 4D Nucleome (4DN) Consortium Omics Standards Working Group and the 4DN Steering Committee (https://www.4dnucleome.org/protocols/). The three human cell lines for data generation were nominated by the 4DN Joint Analysis Working Group and cultured under the 4DN Cell Working Group approved protocols (https://www.4dnucleome.org/cell-lines/). All the data are accessible through the 4DN Data Portal (see Data Availability and Table S1).

The initial challenges to caRNA as a distinct class of RNA were focused on whether these RNAs are exclusively nascent transcripts^9^. Such a concern was alleviated by the discoveries of long-range RNA-chromatin interactions^10–14^, suggesting that caRNA does not completely overlap with nascent transcripts. Our genome-wide analyses reveal two features of RNA-genome association. First, RNA is preferentially associated with its transcription site and up to several megabases of flanking genomic sequence. Second, TAD boundaries insulate RNA-DNA contacts, evidencing the impact of 3D genome on the spatial distribution of caRNA.

It remains unclear how RNA may affect the 3D genome. Because several 3D features of the genome can be reproduced by computational models without considering RNA^64,65^ and *in vitro* experiments to recapitulate loop extrusion without RNA^66^, RNA was not expected to affect the genome’s 3D organization. Furthermore, previous work found that acute reduction of RNA had subtle impacts to the 3D genome at the compartment and the TAD levels ^49^. Our analyses led to similar findings. At the compartment level, Hi-C’s PC1 in FL and RNase exhibited strong correlations with Hi-C’s PC1 in control, suggesting these perturbations had little impact to A/B compartments. At the TAD level, FL and RNase exhibited “highly concordant” TADs with the control, based on the “Measure of Concordance (MoC)”^63^ (pairwise MoCs=0.93 and 0.90, well above 0.75, the threshold for being “highly concordant”^63^). These data confirm that the impacts of RNA to the 3D genome are subtle, at least at the scales of A/B compartments and TADs.

It has not been tested whether an acute reduction of RNA exerts systematic impacts to chromatin loops. Our data reveal either transcription inhibition or acute RNA reduction induced chromatin loops. Most induced loops are shared between transcription inhibition and acute RNA reduction, indicating that the impact on chromatin looping cannot be completely attributed to transcription or the presence of RNAPII on chromatin. Indeed, suppressing a specific caRNA created a chromatin loop, with the loop anchors striding across the genomic sequence associated with this caRNA (Figure 4e-g). Furthermore, deleting the genomic sequence of a caRNA (Chr13:55.5MB_HERV) strengthened the chromatin loops on other chromosomes (Figure 5d). These inter-chromosomal effects argue against that loop strengths are modulated by the transcription of the deleted sequence. They support the idea that the caRNA at specific locations, i.e. between-anchor caRNA, suppresses chromatin looping. Of note, these experiments were not meant to establish an exclusive role of RNA in modulating chromatin looping. While these data establish RNA’s role, they do not exclude transcription or RNAPII’s role in modulating chromatin looping.

What remains to be addressed is whether there is any rule that links specific RNA with specific loops that this RNA can modulate. Disrupting electrostatic interactions by NH4OAc did not lead to significant changes in chromatin loops, withholding us from exploring possible rules based on charge-mediated condensates or phase separation. Instead, we investigated RNA’s target regions, because the genomic locations of the target regions can be compared with the genomic locations of loops. We found a reverse correlation of the caRNA at specific genomic locations, i.e. between loop anchors, and the loop’s strength. With this correlational rule, we hypothesized which RNA suppresses what chromatin loops. We were able to modulate the hypothesized chromatin loops by targeting the identified RNA, thus validating the identified relationships. These validations suggest a general approach based on the localization of caRNA to identify the putative regulatory RNA of chromatin loops.

Recent work revealed a suppressive role of RNAPII’s presence on chromatin to chromatin looping ^42^. Without underestimating RNAPII’s role, our experiments were designed to test if there are any effects of the RNA as well. FL treatment does not displace RNAPII from chromatin ^53^, making the FL emergent loops unlikely due to a change of RNAPII’s presence on chromatin. Furthermore, our analysis focused on the shared loops that are created by both FL or RNase treatment. These shared loops are even more unlikely attributable to the loading of RNAPII on chromatin. Our data suggest that in addition to RNAPII, RNA should be considered toward obtaining a complete picture on the interplay between transcription and genome organization.

## Acknowledgments

We thank 4DN Data Coordination and Integration Center for hosting the raw and the preprocessed datasets for public access. We thank the 4DN Cell Working Group, 4DN Omics Standards Working Group, and 4DN Steering Committee for providing guidelines on experimental protocols and data processing pipelines. We thank the 4DN Joint Analysis Working Group for evaluation of all the data in this resource and cross-comparison with the other 4DN Consortium generated datasets. This work is funded by NIH grants DP1DK126138, R01GM138852, DP1HD087990, and NIH Common Fund 4D Nucleome grants UM1HG011593, UM1HG011585, U54DK107977, and U01CA200147.

## Data availability

All high-throughput data supporting the current study have been deposited on the 4D Nucleome Data Portal (https://data.4dnucleome.org) with the following IDs. iMARGI datasets: H1 control, 4DNESNOJ7HY7; H1 NH4OAc, 4DNESGRI8A8N; H1 FL, 4DNES8B3R3P8; H1 RNase, 4DNESOBRUQ12; HFF, 4DNES9Y1GHK4; K562, 4DNESIKCVASO. Hi-C datasets: H1 control, 4DNESFSCP5L8; H1 NH4OAc, 4DNES2253IBO; H1 FL, 4DNES65I3RQG; H1 RNase, 4DNES4AABNEZ; HFF, 4DNESNMAAN97; K562, 4DNESI7DEJTM.

## Code availability

The codes used for the analysis have been deposited and made publicly available on GitHub at https://github.com/Zhong-Lab-UCSD/RNA3Dgenome-code-repository.

## Extended Text: RBP1 depletion

RBP1, encoded by the POLR2A gene, is the largest subunit of RNA polymerase II. We used the second generation of the auxin-inducible degron (AID2) technology to deplete RBP1^64^. In HCT116 RPB1-Dox-OsTIR1-mClover-mAID cells (RPB1-AID2 cells), where RBP1 is tagged for acute depletion upon addition of doxycycline and 5-Ph-IAA^64^. As the control, we treated RPB1-AID2 cells by doxycycline for 24 hours without 5-Ph-IAA and followed with Hi-C (No-depletion Ctrl), which yielded 632,233,849 read pairs. For the depletion experiment, we treated RPB1-AID2 cells by doxycycline for 24 hours and 5-Ph-IAA for 6 hours and followed with Hi-C (Depletion group), which yielded a comparable number (716,607,191) of read pairs to the IAA-Ctrl. We subjected these data loop calling with HiCCUPS^60^. The Depletion group yielded 3,307 loops, which is approximately 16% more than the detected loops in No-deletion Ctrl (2,619) (p-value = 7.9e-9, paired t-test, chromosome by chromosome). These data suggest depletion of RBP1 led to an increase in loop number.

Taking the union of the loops in the No-depletion Ctrl and the Depletion group, we obtained a total of 5,241 loops (Union loops). We compared the loop strengths (P2LL)^5^ between the No-depletion Ctrl and the Depletion group using all the Union loops. The Depletion group exhibited higher P2LLs than the No-depletion Ctrl (fold change = 1.06, p-value < 2.07e-14, paired t-test), suggesting an increase in loop strengths. Taken together, acute depletion of RBP1 resulted in more and stronger chromatin looping.

## Supplementary material

**Supplementary Figure S1.**
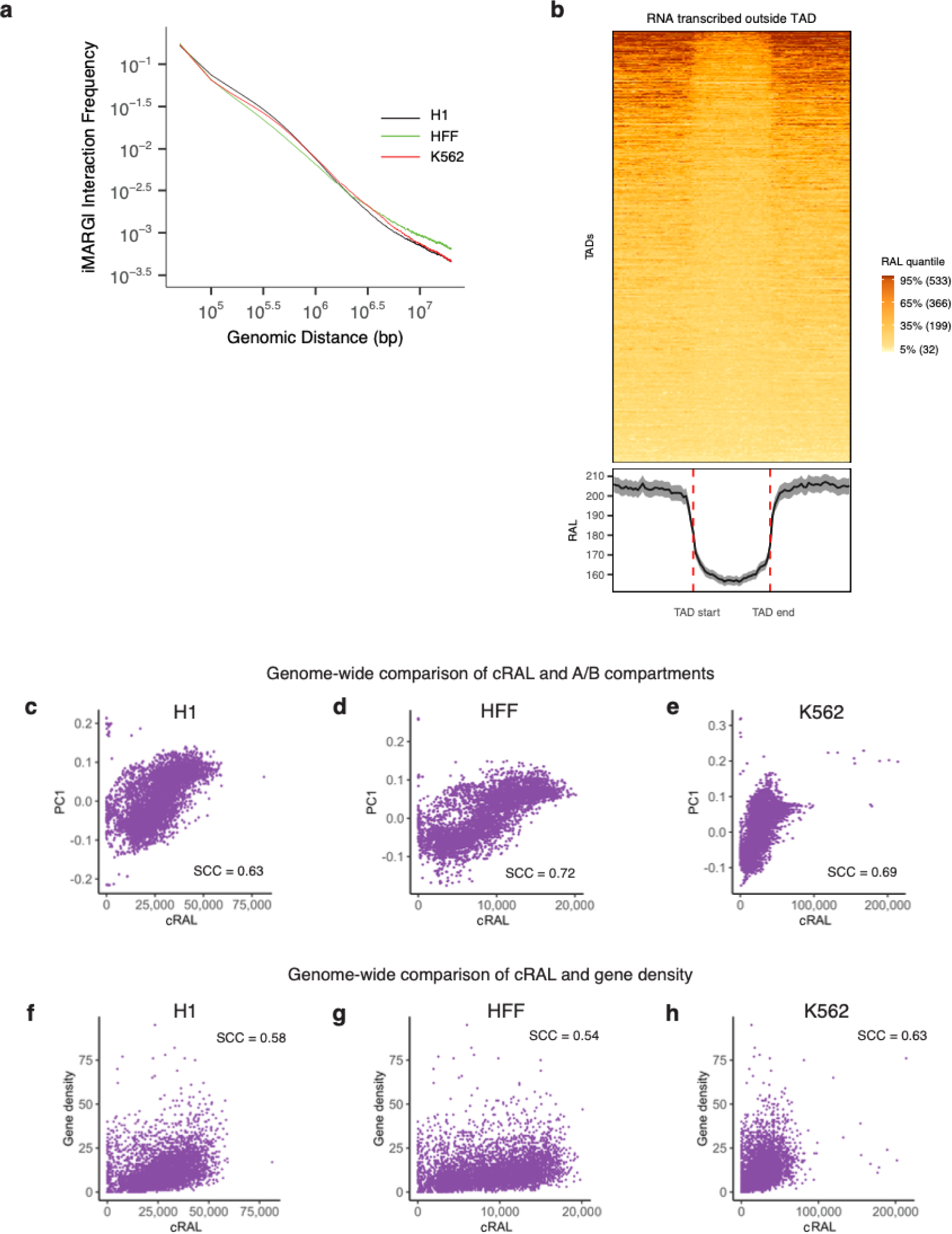
Distribution of RNA attachment levels (RAL) on the genome. (a) RNA-DNA contact ency (y axis) vs. the genomic distance between the mapped RNA-end and the DNA-end (x axis) in H1), HFF (green), and K562 cells (red). (b) The RAL of every TAD (row) and its equal-length flanking regions, based on all the RNAs transcribed from any genomic sequences outside of this TAD (row). The TAD lengths are normalized (center, x axis). Curve at the bottom: average RALs of all TADs. (c-e) Scatterplot of Hi-C’s first vector (PC1, y axis) and cRAL (x axis) on every 500 kb genomic bin (dot) of the entire genome in H1 (c), (d), and K562 (e). SCC: Spearman correlation coefficient. (f-h) Scatterplot of gene density (y axis) and cRAL s) on every 500 kb genomic bin (dot) of the entire genome in H1 (f), HFF (g), and K562 (h). The correlation een cRNA and gene density is weaker than the correlation between caRNA and Hi-C’s PC1.

**Supplementary Figure S2.**
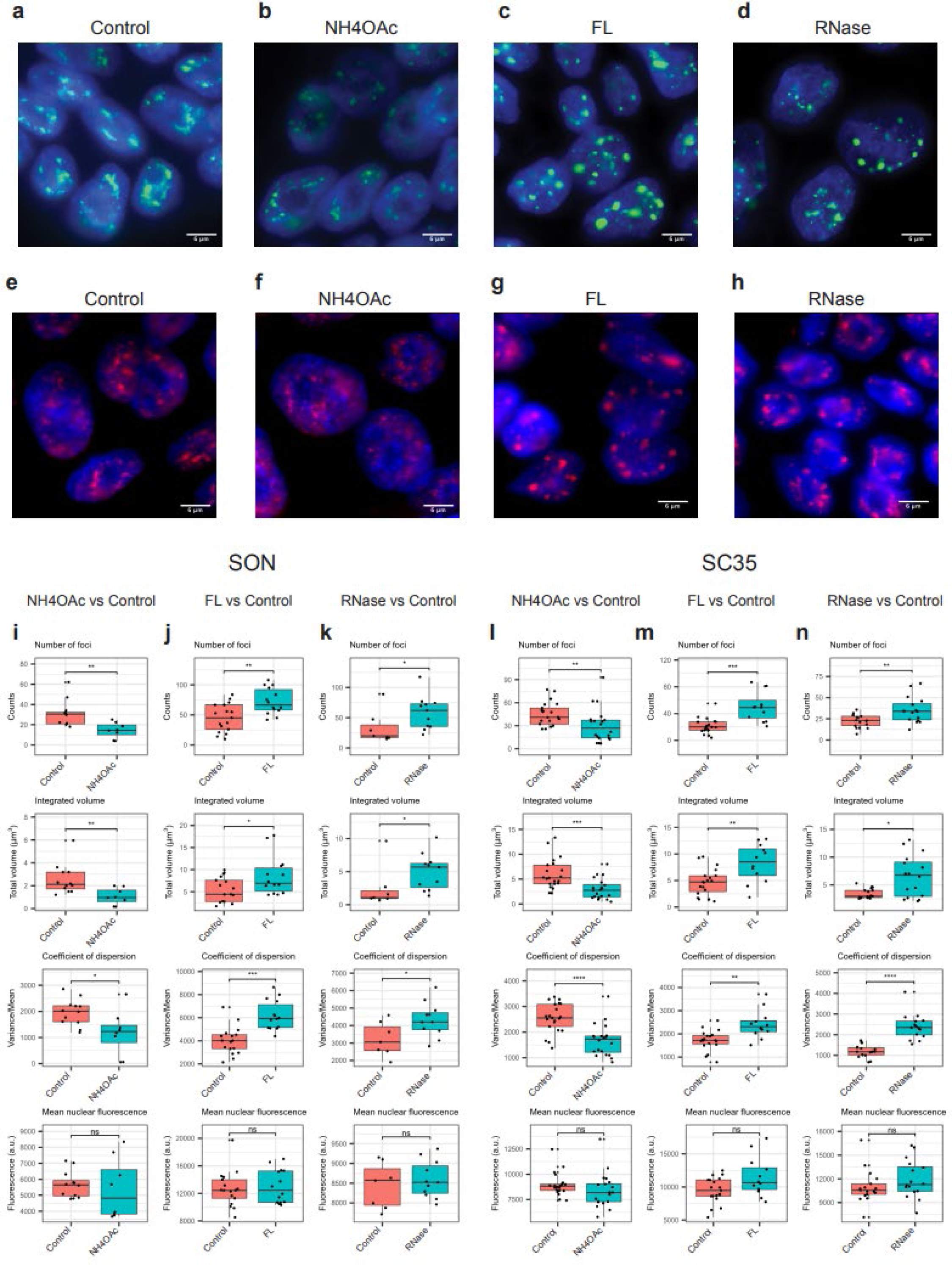
Immunofluorescence analyses of SC35 and SON. (a-h) Immunostaining of SON (a-d SC35 (e-h) in control, NH4OAc, FL, and RNase-treated H1 cells. Scale bar = 6 µm. (i-k) Distribution of s average number of foci per nucleus in control and each treatment (first row). *: p-value < 0.05. **: p-value 1. In comparison, SON’s mean background fluorescence (last row) does not change between control (pink) and each treatment (green). ns: not significant. (l-n) Distribution of SC35’s average number of foci per nucleus in control and each treatment (first row). **: p-value < 0.01. ***: p-value < 1.0e-3. In comparison, SC35’s mean background fluorescence (last row) does not change between control (pink) and each treatment (green). ns: not icant.

**Supplementary Figure S3.**
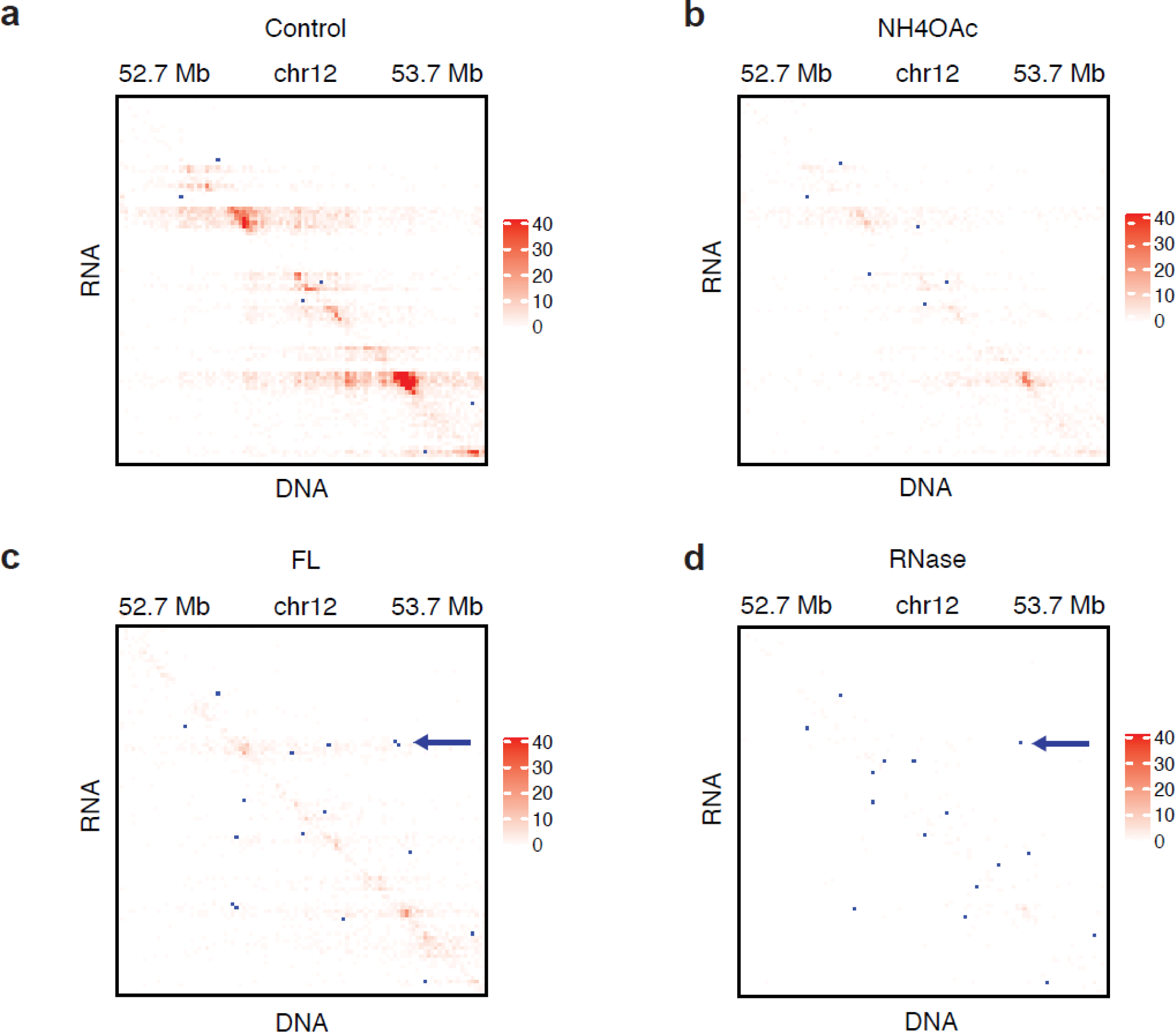
An example of loop change. iMARGI contract matrix of control (a), NH4OAc (b), FL nd RNase (d) for the corresponding genomic region of Figure 3d. Blue dots: Hi-C derived loops that are imposed on the iMARGI contact maps. Arrows: a shared loop in FL and RNase that is absent in control H4OAc.

**Supplementary Figure S4.**
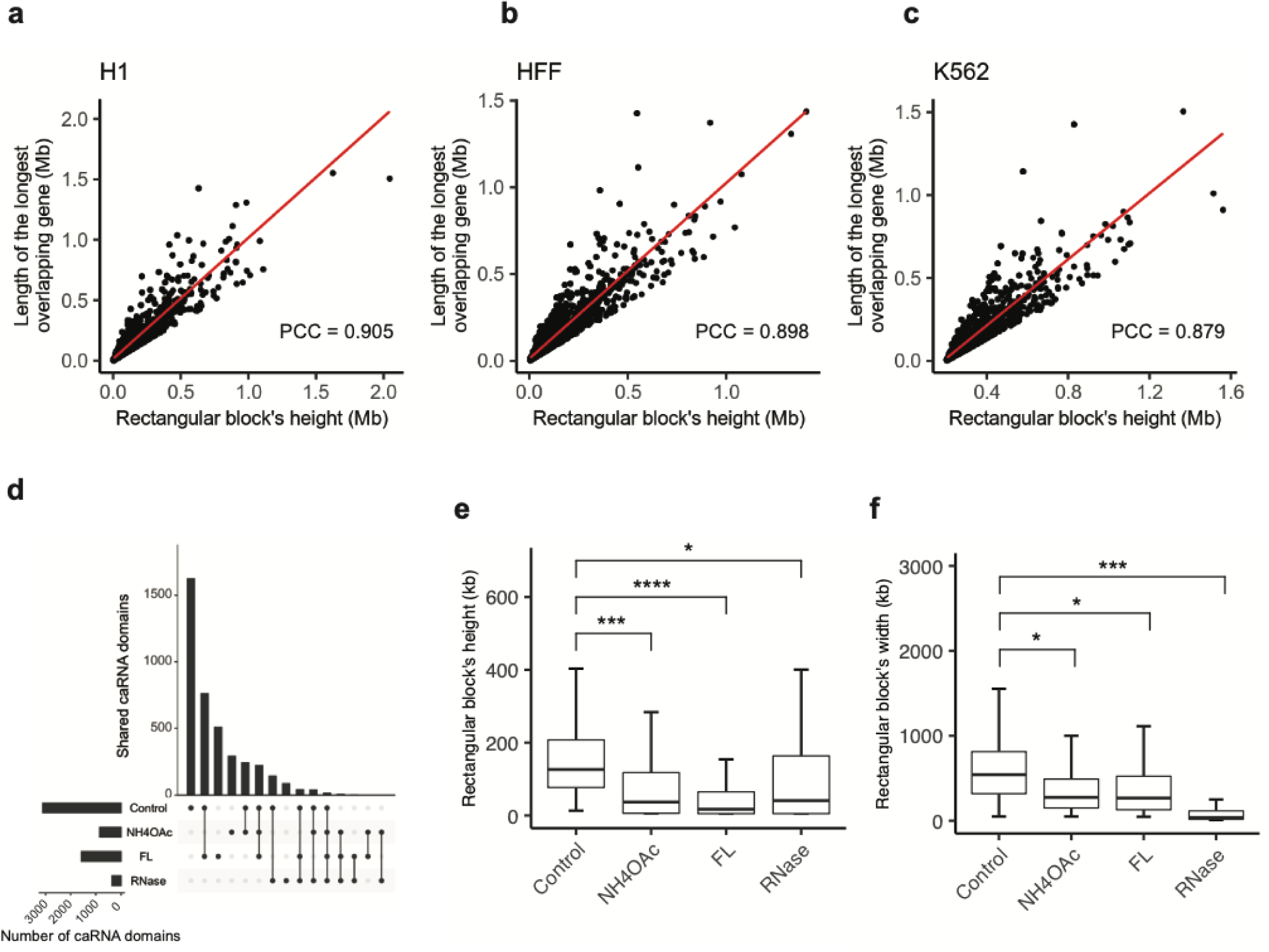
RNA-association domains. (a-c) Scatter plots of each RNA-association domain’s size is) and the length of the longest gene in each RNA-association domain (y axis). Each dot represents an association domain, corresponding to a detected rectangular block from iMARGI’s contact matrix. The and the height of each rectangular block correspond to the size of an RNA-association domain and the h of the genomic sequence that produced the caRNA in this domain. The height of each rectangular block matches the length of the longest gene overlapping with this RNA-association domain, suggesting that RNA-association domains are decorated by the RNA of single genes. (d) Upset plot of the numbers of association domains in untreated H1 (Control) and H1 treated with NH4OAc, FL, and RNase. (e) butions of the heights of the detected blocks, *: p-value < 1e-25, ***: p-value < 1e-75, ****: p-value <1e-100. stributions of the widths of the detected blocks. *: p-value < 1e-50, ***: p-value < 1e-180.

**Table S1.** Summary of iMARGI and Hi-C datasets.

| a. iMARGI datasets |  |  |  |  |
| --- | --- | --- | --- | --- |
| Cell line | Treatment | Number of replicates | Total # of read pairs | Source |
| H1 | None | 4 | 2,642,778,166 | This work |
|  | NH4OAc | 2 | 1,247,204,613 |  |
|  | FL | 2 | 1,438,312,761 |  |
|  | RNase | 2 | 1,670,230,715 |  |
| HFF | None | 2 | 2,755,576,893 |  |
| K562 | None | 2 | 1,293,950,206 |  |
| b. Hi-C datasets |  |  |  |  |
| Cell line | Treatment | Number of replicates | Total # of read pairs | Source |
| H1 | None | 2 | 616,625,628 | This work |
|  | NH4OAc | 2 | 613,098,350 |  |
|  | FL | 2 | 604,503,572 |  |
|  | RNase | 2 | 654,798,738 |  |
| HFF | None | 2 | 2,764,855,452 | 4DNESNMAAN97 |
| K562 | None | 6 | 907,136,828 | 4DNESI7DEJTM |
| c. iMARGI datasets in engineered cells |  |  |  |  |
| Cell line | Treatment | Number of replicates | Total # of read pairs | Source |
| H9 MLC2v:H2B | None | 1 | 692,140,673 | This work |
| H9 MLC2v:H2B<br>HERV2-KO | None | 1 | 566,783,631 |  |
| H9 MLC2v:H2B<br>HERV2-ins-clone2 | None | 1 | 967,136,454 |  |
| d. Hi-C datasets in engineered cells |  |  |  |  |
| Cell line | Treatment | Number of replicates | Total # of read pairs | Source |
| HCT116 RPB1-Dox-OsTIR1-mClover-mAID | Doxycycline<br>(control) | 1 | 632,233,849 | This work |
|  | Doxycycline and 5-Ph-IAA | 1 | 716,607,191 |  |
| H9 MLC2v:H2B | None | 2 | 552,532,207 | GSE116862 |
| H9 MLC2v:H2B<br>HERV2-KO | None | 2 | 635,909,624 |  |

## Online methods

### Cell culture and treatments

Human embryonic stem cells (H1), hTert-immortalized human foreskin fibroblasts (HFF), and chronic myelogenous leukemia lymphoblasts (K562) were obtained from the 4D Nucleome (4DN) Cell Repository and cultured following the 4DN Consortium’s approved culture protocol for each cell line (https://www.4dnucleome.org/cell-lines.html). The cell lines in the 4DN Cell Repository were established by the 4DN Consortium in collaboration with WiCell and ATCC for providing quality-controlled cells from the identical batch to minimize cell source and culture condition variations. The cell culture protocols were developed by the 4DN Cell Line Working Group and approved by the 4DN Steering Committee.

#### Ammonium acetate treatment

H1 cells were treated with 0.1 M NH4OAc in complete mTeSR medium for 10 min as described in a previous study^1^. Briefly, a crystalline NH4OAc (Sigma-Aldrich, Cat# A1542-500G) was dissolved in nuclease-free water and further diluted in cell medium. Aspirate medium in each well and H1 cells were treated with 0.1 M NH4OAc in medium for 10 min at RT.

#### Flavopiridol treatment

H1 cells were treated with 1µM flavopiridol in complete mTeSR medium for 1h in an incubator as described previously^2^. Specifically, a crystalline flavopiridol (hydrochloride) (Cayman Chemical, item# 10009197) was dissolved in DMSO to prepare 1mM flavopiridol (FL) stock solution. 1mM FL stock solution was further diluted with complete mTeSR medium. Aspirate cell medium in each well and H1 cells were either treated with 1µM FL in medium or an equivalent amount of DMSO in the medium in an incubator at 37°C for 1h.

#### RNase A treatment

H1 cells were harvested from cell culture plate and aliquoted cell suspension to 10 million H1 cells per 1.5 mL tube. Wash the cells with 1 mL 1X PBS and centrifuge at 500 X g for 3 min at RT. Then, cells were gently permeabilized by resuspending cell pallets with 0.01% PBST (TritonX-100 in PBS) and treated for 5 min at RT. After permeabilization, cells were treated with 200 µg/mL RNase A as described previously^3^ (Thermo Fisher Scientific, Cat# EN0531) on rotator for 10 min at RT. The treated cells were fixed with 4% formaldehyde (Thermo Fisher Scientific, Cat# 28906) for immunofluorescence imaging. For Hi-C and iMARGI library generation, the treated cells were fixed with 1 mL 1% formaldehyde on rotator for 10 min at RT. Then, the reactions were terminated with 250 µL 1M glycine on rotator for 10 min at RT. The treated sample was centrifuged at 2000Xg for 5 min at 4°C and washed with 1 mL cold 1X PBS.

#### dCas9-KRAB inducible cells

The doxycycline-inducible dCas9-KRAB H1 ES cell line is generated and karyotyped by the 4D Nucleome Consortium (Danwei Huangfu Laboratory) (https://4dnucleome.org), with TRE-dCas9-KRAB and CAGGS-M2rtTA targeted into the AAVS1 locus.

#### HERV-H deletion and insertion cells

The control H9 human ES cells (H9 MLC2v:H2B), HERV-H deletion cell line (H9 MLC2v:H2B HERV2-KO), and HERV-H insertion cell line (H9 MLC2v:H2B HERV2-ins-clone2) were generated by Bing Ren lab and described in reference^4^.

#### RPB1 The auxin-inducible degron 2 cells

The RPB1 auxin-inducible degron 2 cells (HCT116 RPB1-Dox-OsTIR1-mClover-mAID) were generated by Masato Kanemaki lab and described in reference^5^.

### Calling *de novo* TAD boundaries

TAD boundaries in WT (H9 MLC2v:H2B) and KI (H9 MLC2v:H2B HERV2-ins-clone2) were separately called based on their respective Hi-C data using The Arrowhead tool in the Juicer Tools^6^ with default parameters. A TAD boundary called in KI but not in WT is regarded as a *de novo* TAD boundary.

#### Immunofluorescence imaging

The cells on coverslip (Fisher Scientific, Cat# 12-541A) were fixed with 4% formaldehyde at RT for 30 min. The fixed cells were washed with 1X PBS once and permeabilized with 0.1% TritonX-100 in PBS (PBST) at RT for 15 min on shaker. Afterwards, cells were blocked with 5% BSA (VWR, Cat# 97061-420) in PBST at RT for 30 min with gentle shaking. For SC35 staining, H1 cells were incubated with 1 mL diluted mouse monoclonal anti-SC35 primary antibody (1:250) (Abcam, Cat# ab11826) in 5% BSA at 37°C for 1h, and subsequently washed three times with PBST on shaker for 10 min. Cells were further incubated with 1 mL diluted goat anti-mouse secondary antibody with Alexa Fluor 568 (1:500 dilution) (Invitrogen, Cat# A-11004) in 5% BSA at 37°C for 30 min. For SON staining, the cells were incubated with 1 mL diluted rabbit anti-SON primary antibody (1:2000 dilution) (Atlas Antibodies, HPA023535) in 5% BSA at 37°C for 1h, and subsequently washed three times with PBST on shaker for 10 min. The cells were incubated with 1 mL diluted goat anti-rabbit secondary antibody with Alexa Fluor 488 (1:500 dilution) (Invitrogen, Cat# A-11008) in 5% BSA at 37 in 5% BSA at 37°C for 30 min. After staining, the cells were washed three times with PBST on shaker for 10 min. The cells on coverslips were mounted on slides (Fisher Scientific, Cat# 12-544-2) with 10 µL ProLong antifade glass mountant with NucBlue stain (Thermo Fisher Scientific, Cat# P36981), placed in dark room for air-dry overnight. Images in the size of 512×512 pixels were acquired on Applied Precision OMX Super Resolution Microscope using a 100X/1.518 oil objective (GE Healthcare Life Sciences) (pixel size = 0.079 μm). Z-stack images were acquired with thickness of 0.3 μm sample thickness.

#### Identification of nuclear speckle foci

Nuclear speckle foci were identified by a previously described method^2^. Briefly, the nuclei were manually segmented and the mean fluorescence intensity in nuclei were measured with FIJI. The nuclear speckle foci were identified by FIJI 3D Object Counter plugin, with an appropriate intensity threshold of the mean fluorescence intensity in the cell nuclei and a size cut-off of more than 50 adjoining pixels (pixel size, 79 nm X 79 nm).

#### In situ Hi-C library generation and data processing

The Hi-C libraries were generated with the Arima-HiC kit (Arima Genomics, material# A510008, Document# A160134 v00) following the manufacturer’s instructions. Hi-C data was processed following 4DN consortium’s Hi-C data processing protocol (https://www.4dnucleome.org/protocols.html). Briefly, the Hi-C data were processed using the 4D Nucleome (4DN)’s Hi-C Processing Pipeline (v0.2.5) (https://data.4dnucleome.org/resources/data-analysis/hi_c-processing-pipeline), with MAPQ > 30 to filter out multiple mappings.

The output .pairs files were provided to Cooler^7^ (v0.8.10) and Juicer Tools^6^ (v1.22.01) to generate .mcool and .hic files. The .mcool file was used in HiGlass^8^ for visualization. The .hic files were inputted in Juicer Tools for A/B compartment, TAD, and loop analyses. A/B compartments were called by Juicer’s “Eigenvector” tool based on KR normalized observed/expected (O/E) contacts at 500 kb resolution. TADs were called by Juicer’s “Arrowhead” tool based on KR-normalized contacts at 10 kb resolution. Loops were called by Juicer’s “CPU HiCCUPS” tool based on KR-normalized contacts simultaneously at 5 kb and 10 kb resolutions. Except for the resolution parameter, all the other parameters were left as the default.

TAD boundaries were extracted as the genomic regions between TADs in each sample. TAD boundary insulation score was calculated according to the definition in Crane et. al, 2015^9^.

Unique loops and overlapping loops were determined as follows. First, the Juicer called loops from each condition were merged into “unique loops” by taking the union. Then the unique loops in the union were reassigned to each condition by the following rule: a unique loop *i* (in the union) with anchor size *s* (either 5 or 10 kb) was re-assigned to a sample *j* if both anchors of loop *i* were within +/-*s* flanking regions of a loop in sample *j*. Aggregate Peak Analysis was performed using the Juicer’s “APA” tool with default parameters. Metrics to define the loop strength such as Peak to Lower Left (P2LL), Z-score Lower Left (ZscoreLL), and Peak to Mean (P2M) were calculated as defined in Juicer’s APA^6^. The control loop straddling the AAVS1 locus was detected from H1-hESC Micro-C data^10^. To select RNase emergent loops that stride across HERV-H caRNA-attached genomic sequences in Control, i.e., the candidate HERV-H caRNA insulated loops (CHRI-loops), we used a threshold of at least 2 iMARGI read pairs with their RNA ends overlapping with HERV-H and their DNA ends mapped to the between-loop-anchor sequence. To check if deleting a copy of the HERV-H repeats led to increase of loop strengths of CHRI-loops, we used a threshold of 0.05 for the delta peak (KO peak - control peak), where peak is the normalized Hi-C read count at the loop’s pixel, normalized by the total number of read pairs in each sample.

#### Bias correction on Hi-C data for loop visualization

H9 Control and HERV-H2 KO Hi-C data were subjected to HiCorr^11^ with default parameters for bias correction and subsequently subjected to noise removal using the LoopDenoise function in DeepLoop^12^. All data processing was done with Hg19 per HiCorr and DeepLoop software’s requirements.

#### iMARGI library generation and data processing

iMARGI libraries were generated and processed as previously described^13^. According to 4DN’s approved iMARGI’s data processing protocol^13^, any iMARGI read pair in which the RNA end and the DNA end mapped to within 1,000 bp of each other on the genome are removed from the data analysis. The RNA attachment level (RAL) of each genomic segment is the count of the DNA-ends mapped to this genomic segment^14^. Only the inter-chromosomal and the intra-chromosomal iMARGI read pairs that are separated by at least 200 kb apart were used for calculating RAL in any of the correlation analyses. Repeats of hg38 were downloaded from RepeatMasker (Smit, AFA, Hubley, R & Green, P. RepeatMasker Open-4.0). RAL of Alu-containing caRNA (Alu-caRNA) and LINE1-containing caRNA (L1-caRNA) were calculated as the count of the DNA ends mapped to each genomic segment (500 kb size) whose RNA ends mapped to a repeat segment of the Alu or LINE1 family respectively.

#### RNA-defined domains

Each rectangular block on iMARGI’s contact matrix was identified as a peak of the iMARGI’s read pairs’ RNA ends (the height of this RNA peak) and a corresponding DNA peak of the DNA ends (the width of this RNA peak). Homer’s findPeaks function was applied to the RNA ends of iMARGI’ read pairs (peak size = 5,000 bp, minimum peak interval = 12,000 bp) to identify the peaks on the RNA ends (RNA peak). For reach RNA peak, all the iMARGI’s read pairs with their RNA ends inside this RNA peak were retrieved. The retrieved read pairs’ DNA ends were subjected to Homer’s findPeaks (peak size=25,000 bp, minimum peak interval=50,000 bp) to identify the peaks on the DNA ends (DNA peaks). If multiple DNA peaks were reported, the DNA peak with the highest read number was designated as the corresponding DNA peak.

#### Genome coordinates

All plotted genome coordinates are based on Hg38.

